# Compartmentalized cytokine networks and systemic immune remodeling in bovine mammary H5N1 infection

**DOI:** 10.64898/2026.03.06.710145

**Authors:** Gagandeep Singh, Konner Cool, Shristi Ghimire, Jessie D. Trujillo, Igor Morozov, Natasha N. Gaudrault, Huibin Lv, Kaijun Jiang, Yonghai Li, Tuvshintulga Bumduuren, Chester D. McDowell, Adolfo Garcia-Sastre, Nicholas C. Wu, Juergen A. Richt

## Abstract

Highly pathogenic avian influenza A H5N1 has recently expanded its mammalian host range; in 2024, genotype B3.13 emerged in U.S. dairy cattle with pronounced mammary tropism. In the past, Influenza A virus immunology has been characterized primarily in respiratory infection models, whereas this study delineates immune responses after intramammary infection. An intramammary H5N1 challenge in Jersey cows in the early dry-off period enabled integration of dose- and compartment-resolved (alveoli versus teat cistern) cytokine and chemokine profiles with peripheral leukocyte dynamics and H5/N1-specific antibody responses. Infection-induced quarter-restricted, monophasic inflammatory networks peaking at 3–7 days post-infection, coordinated peripheral myeloid expansion and IFN-γ–competent lymphocyte activation, and rising antibody titers across quarters.

## Main text

In March 2024, genotype B3.13 H5N1 viruses were first confirmed in U.S. dairy cattle, with spread across multiple states associated with the widespread detection of viral RNA and high-titer infectious virus in milk, indicating an atypical mammary tropism^1–3^. While influenza A virus infections and their associated immune responses are extensively studied in the respiratory tract, the kinetics, compartmentalization, and cellular correlates of antiviral immunity in bovine mammary tissue remain poorly understood.

A controlled intramammary challenge model in early drying off Jersey cows was performed with the North American HPAI H5N1 isolate A/Cattle/Texas/063224-24-1/2024 (genotype B3.13) to study local and systemic immune responses following infection (detailed methods are provided in the supplementary information)^4^. To increase statistical power while limiting animal use, each mammary quarter received a distinct dose of H5N1 inoculum, permitting within-host dose-response comparisons across compartments. The present study focuses on dose- and time-dependent cytokine and chemokine responses and networks within distinct mammary compartments (alveoli versus teat cistern), characterization of peripheral immune-cell phenotypes, and quantification of virus-specific humoral responses.

Cytokine and chemokine induction was highly localized to the H5N1-infected mammary quarter, with little to no change observed in the mock-inoculated quarter (**Figure 1A**). The magnitude of the immune response depended on both time post-infection (DPI) and inoculum dose, typically peaking during the period of maximal viral replication (approximately 2–7 DPI), consistent with the monophasic kinetics reported in respiratory H5N1 models^5–7^. Notably, the largest cytokine responses were observed in quarters given the intermediate dose inoculum, i.e.10^4^ TCID_50_ per quarter, whereas the lowest dose, 10^3^ TCID_50_ per quarter, produced only minor changes.

**Figure 1.**
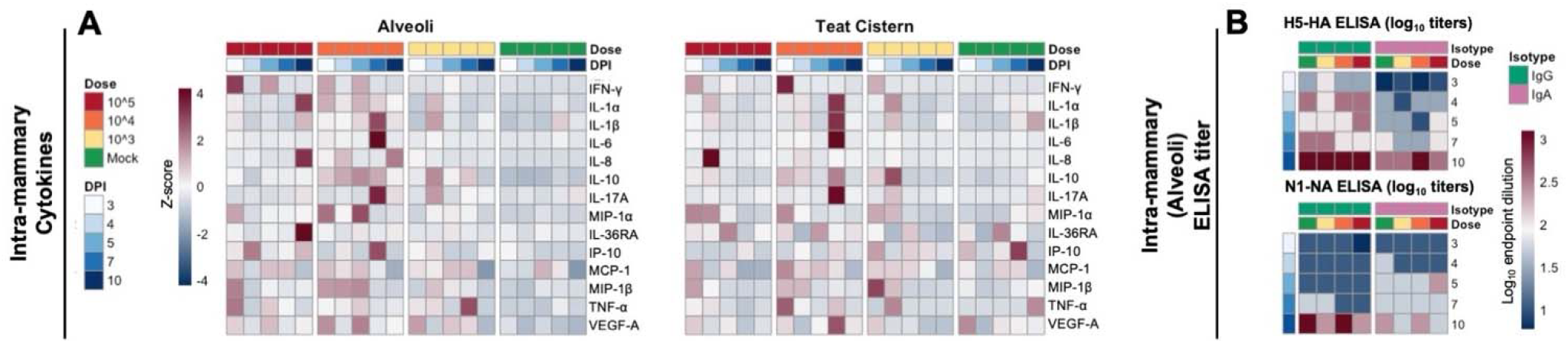
Compartmentalized mammary cytokines and antibodies dynamics in bovine H5N1 infection. (A) Heatmaps of intramammary cytokines/chemokines in alveolar and teat cistern fractions across dose and DPI, showing compartment-specific kinetics. (B) Intramammary H5-HA and N1-NA ELISA titers (log_10_) with IgG/IgA isotype resolution. Number of titer in log_10_.

In alveolar fractions, the intermediate (10^4^ TCID_50_) and high dose (10^5^ TCID_50_) H5N1 inoculations triggered an early induction of IFN-γ, TNF-α, IP-10 (CXCL10), MCP-1 (CCL2), VEGF-A, and MIP-1α/β, consistent with strong chemotactic sensing for monocytes and other myeloid effectors, a pattern described in severe influenza immunopathology^8,9^. This early chemokine and interferon axis was followed by a later induction of classical proinflammatory cytokines (IL-1α, IL-1β, IL-6, IL-8) and cytokines associated with neutrophil and Th17 biology (IL-17A, IL-36RA). In contrast, the low-dose quarters (10^3^ TCID_50_) exhibited only modest, intermediate-level elevations of several cytokines and chemokines (including IL-1α, IL-1β, IL-10, IL-17A, IP-10, MCP-1, TNF-α, VEGF-A, and MIP-1s), indicating a rather blunted and delayed response compared to the mammary quarters inoculated with the higher titer inocula.

Teat-cistern fractions demonstrated a distinct temporal signature, including early IL-36RA induction, highlighting compartment-specific mammary immune regulation and the emergence of discrete inflammatory loci within the mammary gland.

Humoral responses were assessed by quantifying anti–H5 HA and anti–N1 NA antibodies across isotypes, including IgA and IgG in mammary secretions and IgM and IgG in serum **(Figures 1B and 2A)**. A progressive increase across classes over the course of infection is consistent with active B-cell proliferation in the gland and a mucosal-like antibody milieu in which locally produced IgA immunoglobulin is complemented by serum-derived IgG entering the gland via transudation/transcytosis. Unlike the spatially focal cytokine/chemokine responses, antibody increases were not confined to inoculated quarters, with measurable rises also detected in mock-inoculated quarters, suggesting intramammary diffusion across quarters and/or systemic redistribution during acute infection.

Peripheral blood leukocyte populations were profiled by multiparameter flow cytometry, revealing progressive, coordinated remodeling across lymphoid and myeloid compartments **(Figure 2B)**. Within the lymphocyte compartment, IFN-γ–producing cells increased over time, particularly NKp46^+^ NK cells and CD3^+^CD4^+^CD8^+^ T-cell subsets. Myeloid populations were dynamically reconfigured: conventional type 2 dendritic cells (cDC2), CD11b^high^ dendritic cells, and CD80^+^ monocytes expanded substantially, peaking in the early to mid-acute phase, while monocytes and macrophages, including M1-like (CD80^+^CD282^−^) macrophages, remained elevated throughout the entire study period (up to 10□PI). In contrast, plasmacytoid dendritic cells (pDCs) declined and remained reduced during the observed period of infection. These systemic changes reflect the alveolar interferon–chemokine signature: local IFN and chemokine production generates chemotactic cues and activation signals that mobilize monocytes, NK cells, and T-cell subsets from bone marrow and peripheral compartments and drive their functional activation.

**Figure 2.**
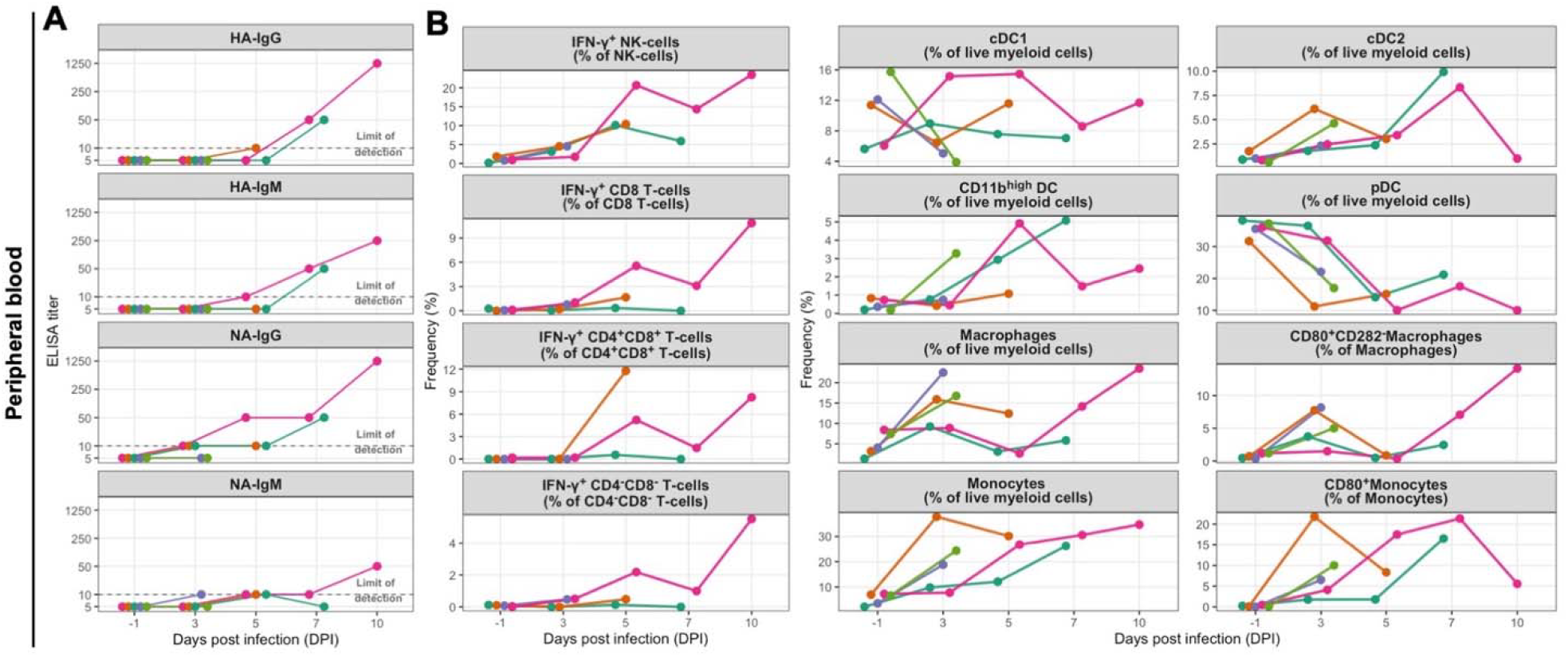
Systemic humoral and immune cells dynamics in bovine H5N1 infection. (A) Peripheral blood HA- and NA-specific IgG/IgM trajectories (Colors represent individual cows). (B) Frequencies of blood IFN-γ^+^ NK/T cells and myeloid/DC/monocyte/macrophage subsets, including activation-marker^+^ populations, across DPI (Colors represent individual cows).

Here, we define compartment- and dose-dependent local and systemic immune responses during bovine intramammary H5N1 infection using a dry-cow challenge model. Inflammatory signaling was largely quarter-restricted due to different virus dose inocula (10^3^, 10^4^ or 10^5^ TCID_50_ per quarter) and differed between mammary compartments (alveoli vs. teat cistern), whereas H5/N1-specific antibody responses increased across all mammary quarters - even mock-infected ones - alongside coordinated changes in circulating myeloid and IFN-γ–producing lymphocyte compartments. These data provide a basis for a better understanding of the mammary-tropic H5N1 pathogenesis in dairy cattle and for linking field surveillance readouts to underlying immune dynamics.

## Acknowledgments

We thank Dr. Diego G. Diel (Department of Population Medicine and Diagnostic Sciences, College of Veterinary Medicine, Cornell University, Ithaca, NY, USA) for providing the HPAI H5N1 isolate A/Cattle/Texas/063224-24-1/2024; Karinne Cortes, Catherine Hickman, Christine Huncovsky, and Michelle Edie for excellent administrative support; and the BRI staff for expert management and operation of the BSL-3 facility.

