## Supplemental Material for "Compartmentalized cytokine networks and systemic immune remodeling in bovine mammary H5N1 infection"

**Material and Methods**

**Ethics statement / Biosafety**

All experiments were approved and performed in compliance with the Kansas State University (KSU) Institutional Biosafety Committee (IBC, Protocol # 1758) and the Institutional Animal Care and Use Committee (IACUC, Protocol # 4992). All animal and laboratory work involving infectious viruses was performed in biosafety level−3+ and −3Ag laboratories and facilities in the Biosecurity Research Institute (BRI) at KSU in Manhattan, KS, USA. The samples were inactivated using approved standard operating procedures, and further evaluation of the inactivated samples was conducted in a BSL-3 laboratory before transferring them to BSL-2 laboratories, where enhanced biosafety practices were employed.

**Cells and Viruses**

Madin–Darby canine kidney (MDCK; ATCC) cells were cultured in Dulbecco’s modified Eagle medium (DMEM; Thermo Fisher Scientific) supplemented with 10% fetal bovine serum (FBS) and 1× antibiotic–antimycotic solution (100 U/mL penicillin, 100 µg/mL streptomycin, and 0.25 µg/mL amphotericin B; Thermo Fisher Scientific). Cells were maintained at 37 °C in a humidified incubator with 5% CO_2_. Sf9 cells (Spodoptera frugiperda ovarian cells; ATCC) were maintained in Sf-900 II SFM medium (Thermo Fisher Scientific) at 28 °C in a shaker incubator.

The highly pathogenic avian influenza virus (HPAIV) strain A/dairy cattle/Kansas/5/2024 was isolated from an HPAIV-affected dairy farm in Kansas and propagated in 9-day-old specific-pathogen-free (SPF) embryonated chicken eggs as previously described^6^. The HPAIV H5N1 isolate A/Cattle/Texas/063224-24-1/2024 was kindly provided by Dr. Diego G. Diel (Department of Population Medicine and Diagnostic Sciences, College of Veterinary Medicine, Cornell University, Ithaca, NY, USA). All viral stocks were titrated by standard plaque assay or TCID_50_ on MDCK cells and stored at −80°C until use.

**Animal experiment design and sample collections**

The samples analyzed in this study were obtained from a parallel experiment currently in preparation. Five female Jersey cows (2–3 years old, monoparous, 700–1,100 lb) in early dry-phase were challenged intramammarily with the North American HPAIV H5N1 isolate A/Cattle/Texas/063224-24-1/2024 (genotype B3.13). Prior to inoculation, the udder was cleaned, and teat orifices were disinfected with 70% isopropyl alcohol or ethanol; virus suspensions (0.5 mL per teat) containing 10^3^, 10^4^, or 10^5^ TCID_50_ were administered into three teats using a teat-infusion cannula, leaving the fourth teat uninfected. After administration, the teat orifices were pinched/held shut for roughly 60 seconds, followed by disinfection with iodine. Blood was collected by jugular venipuncture for immunological assays, and tissues were collected at necropsy on day −1 (pre-challenge) and days 5, 7, and 10 post-challenge (DPC) **(Supplementary Figures 1)**.

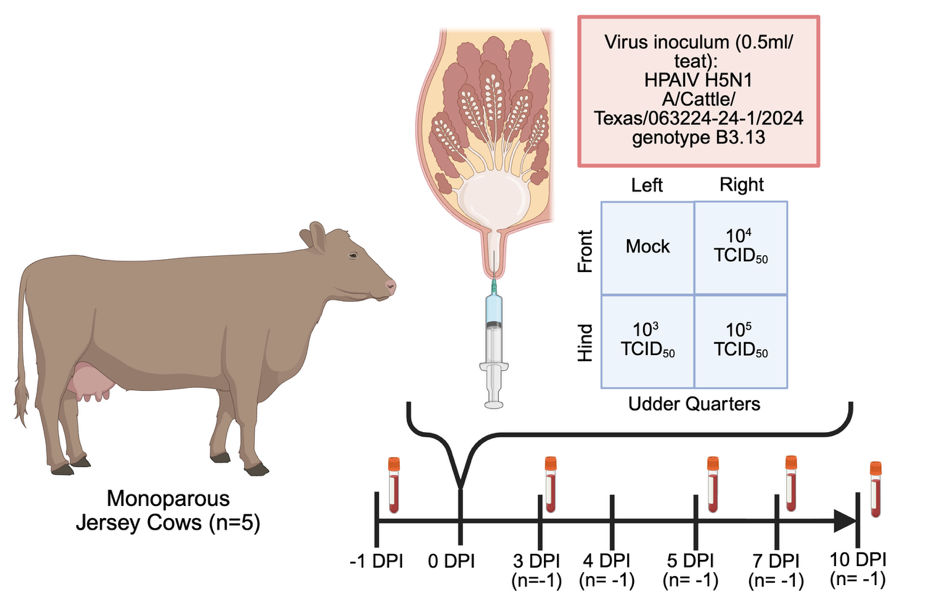

**Supplementary Figure 1. Experimental design and sample collection timeline.**
Schematic representation of the intramammary HPAIV challenge in dairy cows (Created using Biorender).

### **RNA extraction and RT–qPCR analysis for infection confirmation**

To confirm H5N1 infection, clarified mammary gland tissue homogenates (10% weight /volume (w/v)) collected at necropsy on 3, 4, 5, 7, and 10 days post infection (DPI) were mixed 1:1 with RLT lysis buffer (Qiagen) and subjected to total nucleic acid extraction using automated magnetic bead–based platforms (Taco Mini, GeneReach; BioSprint 96, Qiagen) with associated reagents (GeneReach), following established protocols^6^. Extracted nucleic acids were analyzed in duplicate using a one-step RT–qPCR assay targeting the influenza A virus (IAV) matrix gene, employing a modified M + 64 probe and qScript XLT 1-Step RT–qPCR ToughMix (QuantaBio) on a CFX96 Touch Real-Time PCR Detection System (Bio-Rad). Thermocycling conditions were as previously described. Samples were considered positive when both technical replicates yielded cycle threshold (Ct) values ≤38 **(results were presented in the supplementary datasheet).**

**Cytokines and chemokines multiplex analysis**

Clarified mammary gland tissue homogenates (10% w/v) from 3, 4, 5, 7, and 10 DPI necropsy days were used as described above and transferred into UV-transparent U-bottom 96-well plates (Corning). The samples were inactivated with ultraviolet (UV-C) light for 20 minutes with a total UV-C dosage exceeding 2400 mJ/cm^2^. These inactivated samples were then transferred to BSL-2 and used to profile cytokine and chemokine levels using a bovine 15-plex, bead-based Luminex assay (catalogue number BCYT1-33K-PX15, Millipore Sigma). The assay was performed according to the manufacturer’s instructions, with all incubation steps conducted on an orbital shaker set at 250 rpm. Briefly, 25 μl of UV-C-inactivated clarified tissue homogenate supernatant was combined with beads in a low-binding 96-well plate provided in the kit and incubated for 2 hours at room temperature. After incubation, the plate was washed three times with 150 μl/well of wash buffer, followed by the addition of 25 μl/well of the detection antibody mixture for 30 minutes at room temperature. The plate was washed three times, and 25 μl/well of 1× Streptavidin-PE solution was added for 1 hour at room temperature. After three washes, 150 μl/well of Luminex sheath fluid (Millipore Sigma) was added, and the plate was incubated for 5 minutes at room temperature. Data was acquired on a Luminex 200 analyzer (Diasorin) using xPONENT software (version 4.3) and analyzed with R (version 4.4.1) **(see supplementary datasheet for the raw pg/ml value of the cytokines/chemokines)**.

**Expression and purification of HA and NA.**

The HA ectodomain of A/cattle/Texas/56283/2024 (H5N1) was fused with an N-terminal gp67 signal peptide and a C-terminal BirA biotinylation site, thrombin cleavage site, trimerization domain, and a 6×His-tag, and then cloned into a customized baculovirus transfer vector.^1^ The NA head domains of A/cattle/Texas/56283/2024 (H5N1), which contained residues 82 to 469 (N2 numbering), were fused to an N-terminal gp67 signal peptide, 6xHis-tag, a vasodilator-stimulated phosphoprotein (VASP) tetramerization domain, and a thrombin cleavage site. Subsequently, recombinant bacmid DNA was generated using the Bac-to-Bac system (Thermo Fisher Scientific) according to the manufacturer’s instructions. Baculovirus was generated by transfecting the purified bacmid DNA into adherent Sf9 cells using Cellfectin reagent (Thermo Fisher Scientific) according to the manufacturer’s instructions. The baculovirus was further amplified by passaging in adherent Sf9 cells at an MOI of 1. Recombinant protein was expressed by infecting 1 L of Sf9 cells in suspension at an MOI of 1. On day 3 post-infection, Sf9 cells were pelleted by centrifugation at 4000 ×g for 25 min, and soluble recombinant proteins were purified from the supernatant by affinity chromatography using Ni Sepharose excel resin (Cytiva). The protein was further purified by size-exclusion chromatography on a HiLoad 16/100 Superdex 200 prep grade column (Cytiva) equilibrated in 20 mM Tris-HCl (pH 8.0) and 100 mM NaCl. For the NA protein, 10 mM CaCl₂ was included in the buffer. The purified protein was concentrated by an Amicon spin filter (Millipore Sigma) and filtered by 0.22 µm centrifuge tube filters (Costar). The concentration of the protein was determined by nanodrop (Fisher Scientific). Proteins were subsequently aliquoted, flash frozen by dry-ice ethanol mixture, and stored at -80°C until used.

**Humoral immune response against the H5-HA and N1-NA.**

An ELISA targeting the H5-HA and N1-NA proteins was conducted on UV-inactivated serum samples and tissue homogenates. Nunc Maxisorp 96-well plates were coated with 0.1 μg/well (in 50 μL) of recombinant H5-HA or N1-NA and incubated overnight at 4°C using PBS as the coating buffer. The following day, plates were washed three times with PBS-Tween 20 (0.1%) (PBST) and blocked for 2 hours with 2% BSA in PBST at room temperature, followed by three additional washes with PBST. Fifty microliters of five-fold diluted serum (1:5) were added to the wells and incubated for 2 hours at room temperature on a shaker. After three washes with PBST, 100 μL of anti-bovine IgG, IgA, or IgM HRP (horseradish peroxidase) secondary antibodies (Thermo Fisher Scientific) were added to the wells and incubated for 1 hour at room temperature on a shaker. Plates were washed three times with PBST, and a peroxidase reaction was catalyzed using 1-Step TMB (tetramethylbenzidine) ELISA Substrate Solution (Thermo Fisher Scientific). The plates were then incubated for 15 minutes at room temperature in the dark. Finally, the reaction was stopped by adding 50 μL/well of ELISA Stop Solution (Thermo Fisher Scientific). Optical density (OD) readings at 450 nm were obtained using a spectrophotometer (**raw end-point dilutions are presented in the supplementary datasheet)**.

**Peripheral blood collection, PBMC isolation, and lymphocyte stimulation**

Bovine peripheral blood was collected into Na-heparin tubes at −1, 3, 5, 7, and 10 days post-challenge (DPC). Red blood cells were lysed using RBC lysis buffer (8.02 g/L ammonium chloride, 0.84 g/L sodium bicarbonate, 0.37 g/L EDTA), and leukocytes were washed twice in FACS buffer (1× PBS, 0.5% BSA, 0.5 mM EDTA) to isolate peripheral blood mononuclear cells (PBMCs).

For lymphocyte stimulation, PBMCs were seeded at 1 × 10^5^ live cells/well in tissue-culture-treated 96-well plates in RPMI supplemented with 10% fetal bovine serum, 1× 2-mercaptoethanol, and 1× antibiotic–antimycotic (Thermo Fisher Scientific). Cells were inoculated with 1 × 10^5^ PFU/well of bovine H5N1 HPAIV (A/dairy cattle/Kansas/5/2024) and incubated for 4 h at 37 °C in a humidified 5% CO₂ atmosphere. A protein transport inhibitor (Invitrogen) was then added, and incubation continued until a total stimulation time of 24 h was reached. After stimulation, cells were washed and processed for surface and intracellular staining as described below.

Isolated PBMCs and stimulated cells were blocked in FACS buffer supplemented with 2% FBS for 5 min at room temperature, then stained for 30 min at room temperature in the dark with the following surface markers: anti-CD3 (CD3-12), CD4 (CC8), CD8 (CC63), NKp46 (AKS1), CD9 (MM2/57), CD11b (MM12A), CD11c (BAQ153A), CD80 (IL-A159) and CD282 (AbD12542). Cells were washed twice with FACS buffer, fixed in 10% formaldehyde at 4 °C for 4 h, and permeabilized with 1× permeabilization buffer (Invitrogen) for 1 h. Intracellular staining for IFN-γ was performed for 1 h using an anti-bovine IFN-γ antibody (CC302). After two final washes in FACS buffer, samples were filtered through a 100 µm cell strainer (Falcon) to remove clumps and analysed on an Attune NxT Flow Cytometer (Thermo Fisher Scientific). Data were processed in FlowJo v10.10.0 (BD) (**Supplementary Fig. 2, frequencies of the cells are presented in the supplementary datasheet**).

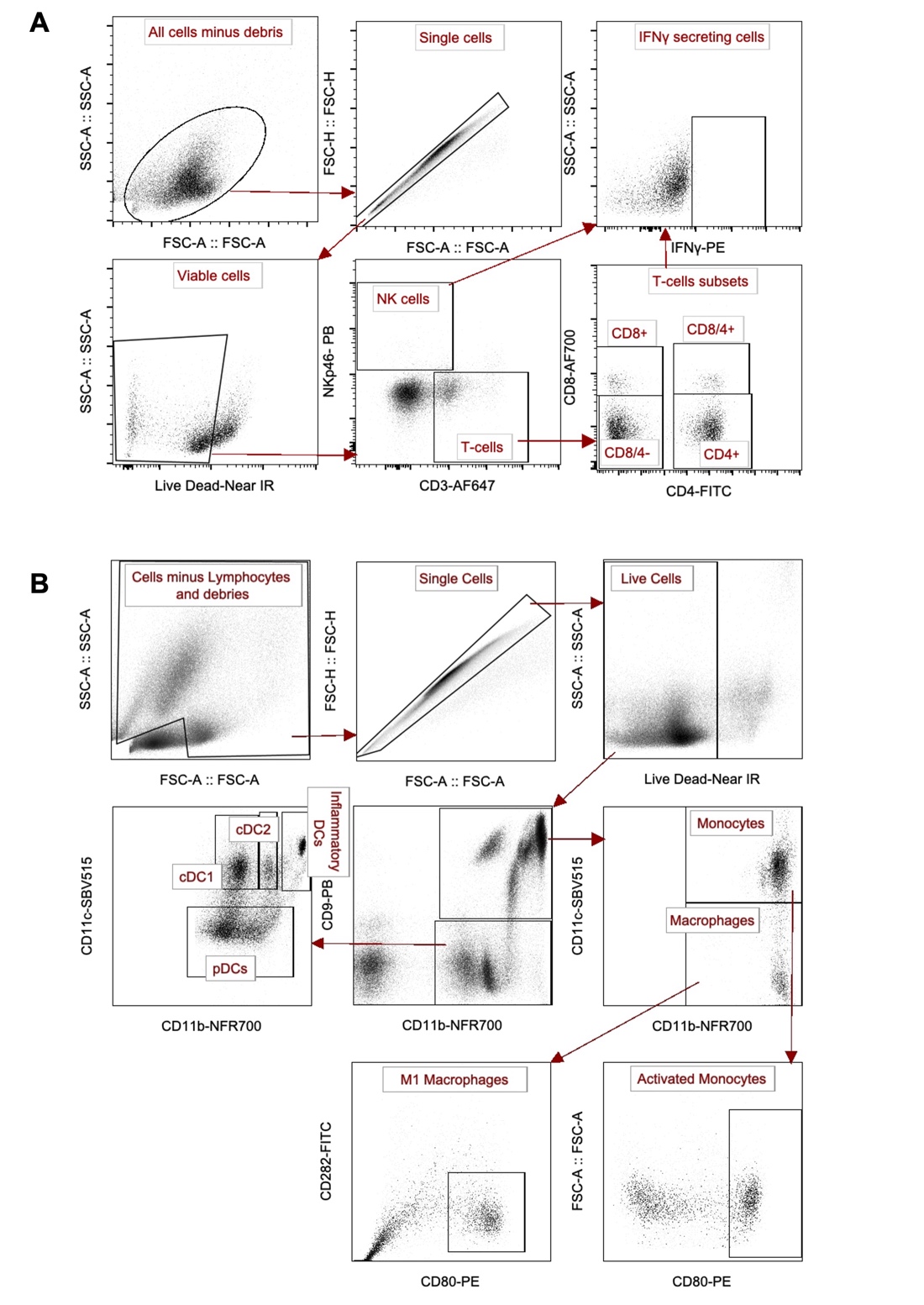

**Supplementary Figure 2. Gating scheme for peripheral blood leukocyte populations profiling.** (A) Lymphocyte compartment (B) Myeloid compartment.
